# NEMAT: An Automated Non-Equilibrium Free-Energy Framework for Predicting Ligand Affinity in Membrane Proteins

**DOI:** 10.64898/2025.12.09.692605

**Authors:** Albert Ortega-Bartolomé, Ramon Crehuet

## Abstract

Quantifying the strength of small-molecule binding to proteins is essential for understanding biological function and for advancing drug discovery. Computational free-energy methods aim to predict these interactions accurately, yet membrane proteins remain particularly difficult due to the added thermodynamic effects of the lipid bilayer. Alchemical free-energy calculations provide state-of-the-art accuracy by transforming one ligand into another, and recent non-equilibrium approaches (NEQ-FEP) offer improved efficiency and parallelization. However, no existing workflow automates NEQ-FEP for membrane proteins and enables decomposition of the observed binding free energy into membrane-partitioning and protein-specific components. NEMAT is an open-source pipeline that performs automated non-equilibrium alchemical transformations in water, membranes, and membrane-embedded protein environments. Here we show that NEMAT reproduces experimental binding-energy trends for P2Y_1_ ligands with accuracy comparable to established equilibrium methods. These findings demonstrate that NEQ-FEP can be applied reliably to membrane systems when supported by a consistent workflow. In contrast to previous approaches, NEMAT provides systematic control over transition length, sampling, and replica averaging, enabling stable work-distribution overlap and reproducible free-energy estimates. In a broader context, NEMAT offers a practical route to mechanistically interpretable affinity predictions for membrane-embedded targets. Its ability to dissect membrane and protein contributions advances our understanding of ligand selectivity at lipid-facing pockets and supports the routine application of non-equilibrium free-energy methods in drug discovery.

## Introduction

Accurately determining the binding free energy (BFE) of protein–ligand complexes remains a central challenge in computational chemistry, with major implications for drug design, and chemical biology in general (1). Despite decades of progress, predicting BFEs is still difficult because of the complex balance between enthalpic and entropic effects and the need for extensive sampling (2). Even long molecular dynamics simulations often fail to produce reliable or reproducible binding free energies (3).

Alchemical transformation (AT) methods offer one of the most accurate strategies for computing relative binding free energies (RBFE) (4), i.e., the difference in BFE of two ligands or two protein variants. These methods estimate RBFE by gradually “mutating” one molecular system into another through a nonphysical —or “alchemical”— pathway. Using a coupling parameter *λ* ∈ [0, 1], which represents the original and modified molecular systems at *λ* = 0 and *λ* = 1, respectively, AT interpolates between molecular Hamiltonians to calculate RBFE, solvation energies, or mutational effects with high precision (5, 6). AT methods are particularly useful for comparing ligand affinities for the same receptor or for studying point mutations in a protein–ligand complex.

Several rigorous statistical mechanics techniques implement alchemical transformations, including Free Energy Perturbation (FEP) (7), Thermodynamic Integration (TI) (8, 9), and non-equilibrium approaches based on Crooks’ fluctuation theorem (10) and Jarzynski’s equality (11). Alchemical FEP methods reduce cost, accelerate results, and improve insight into binding mechanisms and lead optimization (12–14). Consequently, several workflows now automate these approaches (4, 15–18). TI and equilibrium FEP remain widely used for their strong theoretical foundation and consistent performance, but non-equilibrium FEP (NEQ-FEP) has recently become a robust alternative (19, 20). NEQ-FEP offers similar accuracy with better parallelization and efficiency, especially when combined with estimators such as Bennett’s Acceptance Ratio (BAR) (21). In 2019, Gapsys *et al*. (12, 22) released a new NEQ-FEP implementation using GROMACS (23) and pmx (24), making the approach more accessible and reliable.

Despite these advances, alchemical methods remain underused for membrane proteins, particularly with NEQ-FEP (20). As membrane proteins represent many drug targets and are central to key biological processes and diseases (25), the lack of methods to calculate their RBFE with different ligands hinders progress in medicinal and biological chemistry. Dickson *et al*. studied GPCRs with membrane-exposed binding pockets (26), while Zhang and Im examined GPCRs with internal pockets (27). Both studies, performed with TI and AMBER, showed that membranes strongly influence binding energetics when pockets contact the lipid environment. Explicitly including membranes is therefore essential, since ligands may appear to bind tightly merely because they partition into the bilayer rather than engaging in specific protein–ligand interactions (28–30). Partitioning can also increase local ligand concentration near the receptor (31). Additionally, membrane components such as cholesterol can stabilize proteins or modulate binding (32, 33). Distinguishing membrane partitioning from specific binding is thus critical for understanding mechanisms and improving ligand selectivity. Including ligand partitioning in free energy calculations allows decomposition of the observed binding energy (Δ*G*_obs_) into membrane insertion (Δ*G*_mem_) and specific interaction (Δ*G*_int_) terms, as illustrated in Fig. 2.

**Fig. 1.**
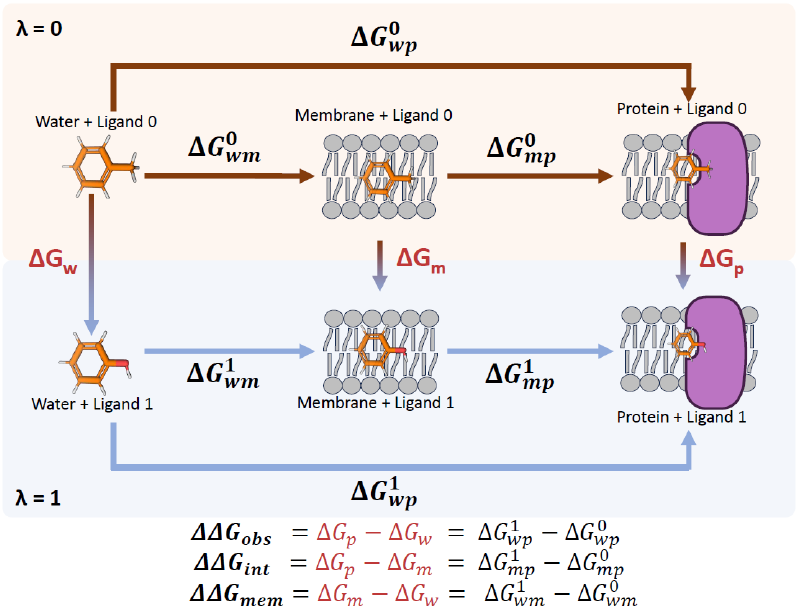
Cycles involved in a NEMAT run. Alchemical transformations consist of transforming ligand 0 to ligand 1 in water, membrane, and protein systems. In the image, ATs are colored in red. Below are the corresponding equations for every part of the cycle. Only the horizontal arrows will be simulated.

**Fig. 2.**
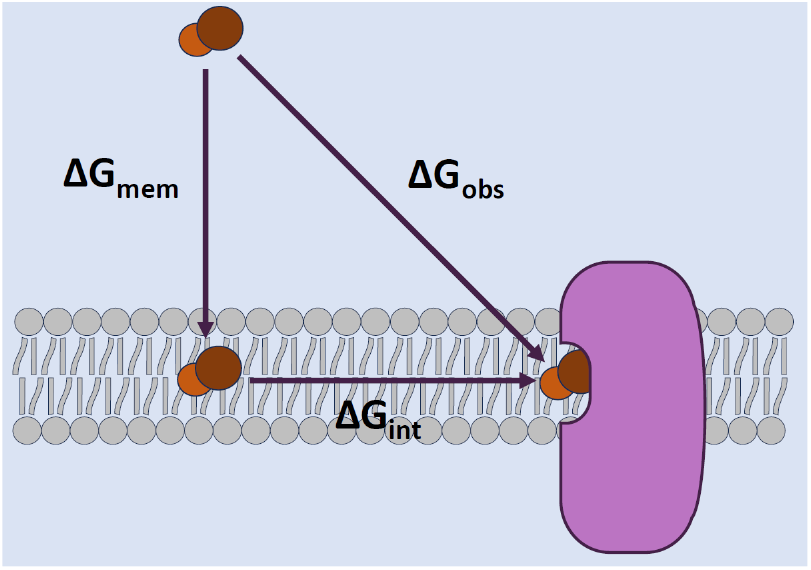
Definition of the Different Binding Free Energies. The binding free energy of a ligand to a membrane protein (Δ*G*_obs_) can be decomposed into two steps: the ligand binding to the protein (Δ*G*_mem_) plus the binding from the membrane to the protein (Δ*G*_int_).

To our knowledge, only one study has explicitly decomposed Δ*G*_obs_ using TI (26), and no established software or protocol supports this approach for NEQ-FEP. To address this gap, we developed NEMAT (Non-Equilibrium Membrane Alchemical Transformations), an open-source pipeline that automates setup, execution, and analysis of RBFE calculations for small molecules in three systems: water, membrane, and membrane-embedded protein (MEP) (Fig. 1).

### A.A practical case: P2Y_1_

To evaluate NEMAT and illustrate its practical use, we reproduced the benchmarks originally performed by Dickson *et al*. using the P2Y_1_ receptor, a class A GPCR. P2Y_1_ is a well-established model for studying ligand binding at membrane-exposed sites, where accurate treatment of both membrane partitioning and protein interactions is essential (26). This receptor binds several lipophilic antagonists, including BPTU, within a lipid-facing allosteric pocket located between transmembrane helices I–III rather than at the conventional orthosteric site (34).

The BPTU chemotype and its series of 30 analogues provide an ideal test set for assessing relative binding free energy (RBFE) methods in membrane environments since these ligands must first partition into the lipid bilayer before engaging the receptor.

Using this system as a benchmark, therefore, serves two purposes: it demonstrates how NEMAT is applied in practice, and it enables systematic evaluation of which simulation parameters most strongly affect accuracy in membrane-protein RBFE calculations.

This work introduces NEMAT, a software for performing AT calculations on membrane proteins, using non-equilibrium methods. We evaluate optimal parameter choices, and compare its performance against a previous equilibrium AT benchmark. We also provide a user tutorial and highlight the importance of accounting for membrane affinities when assessing ligand selectivity for membrane-embedded binding sites. The software repository from NEMAT, with the tutorial files and the code to run it for any new protein, can be found in the NEMAT GitHub repository.

## Methods

### A. The NEMAT pipeline

NEMAT is a Python3/Bash program that automates non-equilibrium free-energy perturbation (NEQ-FEP) calculations in water, lipid bilayers, and membrane-embedded protein (MEP) environments. Its workflow, based on the tutorial code for FEP using pmx (35) (Fig. 3), combines pmx for system setup and analysis with GPU-accelerated GROMACS simulations. Ligand topologies are generated through ACPYPE using the GAFF2 force field, which has been tested for NEQ methods (36). Although CHARMM-GUI (37) is recommended for preparing protein and membrane systems, NEMAT accepts any valid GROMACS-ready membrane–protein setup. It balances flexibility and fine-grained control with ease of use.

**Fig. 3.**
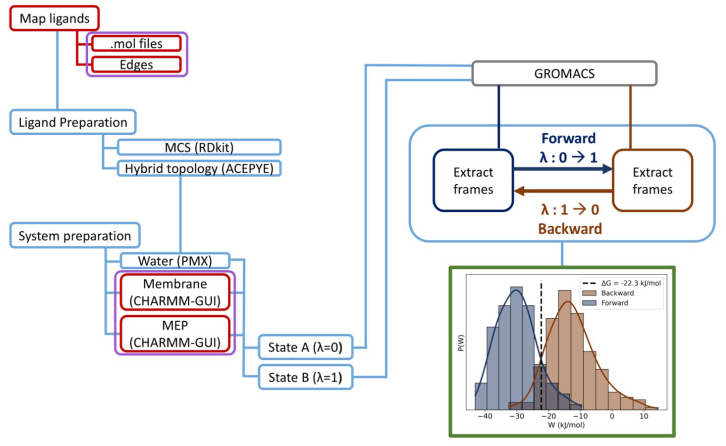
Simple schematics of the NEMAT usual workflow. Red labels highlight the actions the user performs, purple labels indicate the inputs provided to NEMAT, blue elements show the operations executed by NEMAT, and green elements represent the program’s outputs.

- Generation of hybrid topologies via maximum common substructure (MCS) mapping.
- Setup of alchemical transformations across chosen edges.
- Execution of NEQ-FEP simulations in parallel replicas.
- Analysis of relative binding free energies (ΔΔ*G*).

pmx prepares ligands for alchemical transformations and has demonstrated state-of-the-art accuracy in published bench-marking studies using open-source tools (12, 38). It performs especially well for small to medium-sized protein–ligand systems.

### B. System Preparation

Following Fig. 3, users first prepare the MEP, membrane, and ligands. NEMAT begins from pre-built systems without ligands; CHARMM-GUI is recommended for preparing MEP and membrane systems (37, 39–41), though any well-defined GROMACS coordinates and topology files are accepted.

Ligands are provided as .*mol2* files. NEMAT automatically selects alchemical atoms through a maximum common substructure (MCS) search between ligand pairs using pmx. Shared atoms remain across the alchemical pathway, while unique atoms become dummy atoms that disappear or appear along the *λ*-interpolation. This ensures consistent hybrid topologies with both real and dummy atoms for smooth free-energy transitions (24, 42). pmx relies on ACPYPE (43) to supply force-field parameters. ACPYPE wraps Antechamber to generate GROMACS-compatible topologies for small molecules under GAFF2. pmx thus provides atom mapping and hybrid-topology generation, while ACPYPE ensures chemically accurate parameters before alchemical simulations.

### C. Running NEMAT

After system preparation, NEMAT performs free-energy simulations for the three environments—water, membrane, and membrane-embedded protein (MEP)—using the GROMACS software. Following the thermodynamic cycles in Fig. 1, relative free energies are derived: ΔΔ*G*_obs_ (from MEP and water), ΔΔ*G*_int_ (from membrane and MEP), and ΔΔ*G*_mem_ (from membrane and water).

To ensure robust statistics, more than one replica (default is 3) is recommended for each system. Each simulation begins with energy minimization, followed by NPT restrained equilibration. For MEP and membrane systems, equilibration proceeds in six stages with progressively weaker restraints, ending with an unrestrained equilibration to stabilize lipids (44).

After equilibration, an NPT production trajectory is run from which a series of alchemical transformations (from now on, transitions, since the transformation is a transition from a state A (ligand 1) to a state B (ligand 2)) are used to compute the RBFE, as done in NEQ-FEP (45) (Fig. 4). The recommended length of this trajectory will be discussed in the results section. Reported relative binding free energies (RBFEs) are averaged across replicas.

**Fig. 4.**
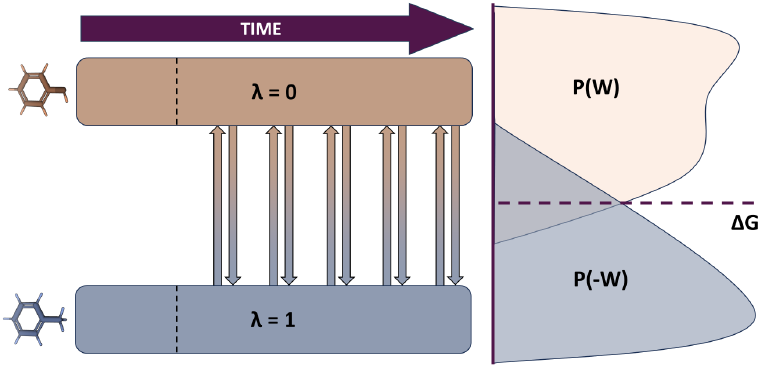
Free energy calculation with Non-equilibrium free energy perturbation. The Molecular Dynamics trajectories for the two ligands (*λ* = 0 and *λ* = 1) are split into an equilibration phase (before the dashed line) and a production phase. From the production phases, fast non-equilibrium transitions are performed forward (from 0 to 1) and backwards (from 1 to 0). The free energy is obtained from the work distribution of the transitions using different estimators.

Because NEMAT uses NEQ-FEP, transitions must remain short to capture the instantaneous work changes during ligand swapping. Several estimators can produce the free energy difference from the work distribution. The simplest and seminal estimator is the Jarzynski equality (Eq. 1, (11))

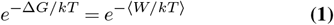

However, the Bennett Acceptance Ratio (BAR) method (21) provides a more reliable estimator by optimally weighting forward and reverse work distributions (46). NEMAT calculates free energies using Jarzynski’s equality, its Gaussian approximation, and BAR, but reports only BAR-derived results due to their superior convergence when bidirectional sampling is available.

Average work values derive from overlapping forward and reverse work distributions (4); adequate overlap is essential for numerical stability. Following best practices (47), users should calibrate parameters per target rather than relying on fixed settings. We recommend validating a small subset of edges first, tuning transition durations and parameters until they meet overlap/uncertainty criteria (ideally, target ≈0.1 overlap score on the MEP system).

When multiple replicas are run, NEMAT computes a Gaussian-weighted average for each system. The replica weight (*ω*_*i*_) is given by:

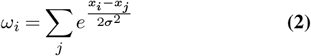

Where *σ* (which is set to 1) controls the relevance of having closer weights. Then, the values of Δ*G*_env_ (representing the free energy difference marked in red in Fig. 1 of the alchemical transformation from ligand A to ligand B in the same system) are computed with the following average:

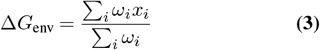

Where *i* is the replica number and env ∈ [w, m, p].

### D.A case example: P2Y_1_ with BPTU analogs

We used the same 14 antagonist ligands tested by Dickson et al. (26), each with measurable *K*_*i*_ values for P2Y_1_. Chao et al. (48) determined these affinities using radioligand binding assays, from which Δ*G* was derived as Δ*G* = −*RT* ln(*K*_*i*_) at 298.15 K. This Δ*G* corresponds to Δ*G*_obs_.

The P2Y_1_–BPTU complex (4xnv, 2.2 Å resolution (34)) was downloaded from the OPM database (49). Following Dickson *et al*., we selected 14 ligands (out of 29 candidates), divided them into two groups (11a-like BPTU analogues and 6a-like BPTU analogues, Fig. S6, S7), and aligned them with the BPTU reference using RDKit.

#### D.1. Membrane-embedded protein (MEP) system: the P2Y1-POPC membrane complex

The P2Y_1_ protein was protonated at pH 7.4. The OPM structure lacks ICL3 due to disorder; however, this region does not affect ligand binding, so the missing residues were not modeled. Termini were capped with ACE and NME. Amber ff19SB was used for the protein (50). A POPC (1-Palmitoyl-2-oleoyl-sn-glycero-3-phosphocholine) bilayer (9.5 x 9.5 nm, 233 lipids) was constructed after aligning the protein along the Z-axis. Lipid parameters used Lipid21 (51). The system contained 0.15 M NaCl (56 Na^+^, 76 Cl^−^) and 20,939 TIP3P water molecules (52). Waters inside the helical bundle were added with Dowser++ (53) to improve realism. Fig. 5 shows the final complex.

**Fig. 5.**
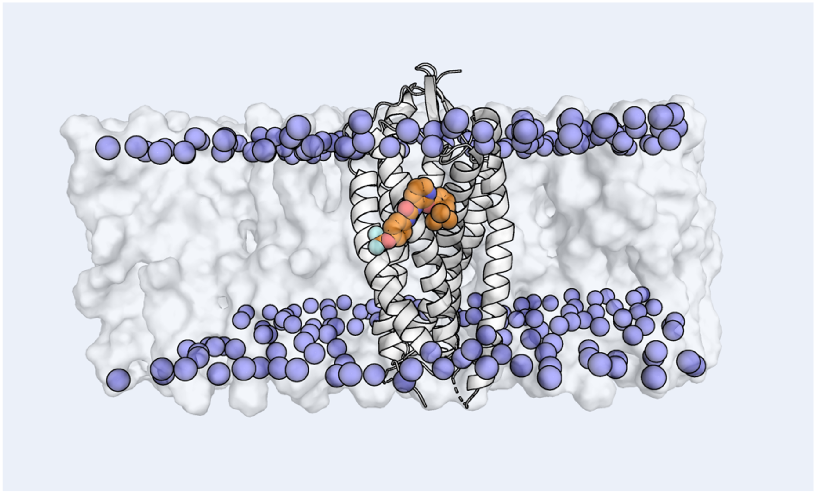
Membrane-Embedded P2Y_1_ Receptor. Representation of the equilibrated membrane-embedded P2Y_1_ receptor including the BPTU ligand (orange) in the binding pocket. Water molecules and ions are not expressed for clarity. The polar heads of the lipids in the bilayer are represented in blue.

#### D.2. Membrane system

A POPC bilayer (4 × 4 nm, 48 lipids) was built with 0.15 M NaCl (5 ions each type) and 2,132 TIP3P water molecules.

### D.3 Small molecules: BPTU analogues

The P2Y_1_–BPTU complex was aligned with the P2Y_1_-POPC membrane complex, and the ligand was extracted as a reference for FEP alignment. As Dickson *et al*. already defined the candidate ligands (supplementary Fig. S6, S7), that selection step was omitted. Ligand topologies and GAFF2 parameters with AM1-BCC charges were generated using ACPYPE. Each ligand was solvated in a 4.5 nm cubic box with ≈ 1,990 water molecules and ionized with NaCl (six ions per type for neutral ligands). Then, pmx performs alchemical mapping.

#### D.4. Running NEMAT

All free-energy simulations were performed with GROMACS 2024.2 using the NEMAT NEQ-FEP protocol.

Three replicas were run to improve convergence. All systems underwent a 5,000-step energy minimization (tolerance 1,000) via steepest descent, followed by a NPT equilibration of 1.5 ns for the systems containing membrane and 0.5 ns for the ligands in water, using the C-rescale barostat. For membrane and MEP systems, equilibration included six stages with progressively weaker restraints (1,000 → 100 kcal/mol/Å^2^ for bilayer; 4,000 → 50 kcal/mol/Å^2^ for MEP) and a final unrestrained phase.

After equilibration, 20 ns NPT production runs were performed, saving 200 frames at regular intervals. Starting from 5 ns, 50 evenly spaced alchemical transitions were executed with 100 ps extensions, using a 2 fs timestep and stochastic Langevin dynamics. These transitions were implemented via GROMACS free-energy options, and all simulations used identical protocols and parameters.

## Results and Discussion

### A. Performance of NEMAT on the P2Y_1_–BPTU Ligand Series

We evaluated NEMAT on the set of BPTU analogues previously characterized for antagonistic activity at the P2Y_1_ receptor. Correct prediction of the sign and magnitude of relative binding free energies (ΔΔ*G*_obs_) is critical for guiding lead optimization, as negative values indicate improved affinity and positive values indicate decreased binding.

We used the experimental ΔΔ*G*_obs_ values reported by Wang *et al*.(54) and curated by Zhang *et al*. (34) to benchmark NEMAT. In addition, we compared the results with an Amber-based equilibrium free-energy perturbation study performed by Dickson *et al*. (26).

For the 11a-centered star map, NEMAT correctly reproduces the experimentally observed ranking of ligands relative to 11a (Tab. 1). All predicted ΔΔ*G*_obs_ values show the correct direction of affinity change, indicating that the workflow reliably distinguishes affinity gains from losses. We observe a similar trend for the 6a-centered series (Tab. 2), with correct directional predictions in 87.5% of ligand pairs (taking into account that the experimental values for *6a*→ *6f* and *6a* →*6j* are essentially 0). These results demonstrate that NEMAT provides accurate prioritization guidance for structure–activity relationship (SAR) exploration.

Beyond predicting bound-state affinities, NEMAT also estimates ΔΔ*G*_int_, the free-energy difference between the membrane and the receptor-bound states. This metric separates true improvements in receptor engagement from changes driven primarily by altered membrane partitioning. In contrast to the values computed by Dickson *et al*., NEMAT produced ΔΔ*G*_int_ estimates that did not follow the same trends, suggesting methodological differences between the two approaches rather than disagreement with experimental data.

NEMAT reproduced the experimental trends with an RMSE of 1.74 kcal·mol^−1^. Its agreement with the values reported by Dickson *et al*. was similarly strong, with NEMAT deviating from their predictions by an average of 1.83 kcal·mol^−1^.

Therefore, NEMAT achieves predictive accuracy comparable to state-of-the-art alchemical workflows while incorporating NEQ-FEP, membrane systems, fewer user-defined steps, and maintaining full reproducibility through its automated pipeline.

### B. Computational Efficiency and Parallelization

Because alchemical transformations in each environment (water, membrane, and MEP) are independent across systems and replicas, the NEMAT workflow is highly parallelizable. When all replicas were executed simultaneously, the total wall time per edge was determined primarily by the MEP system, which was the largest. This parallel structure permitted rapid exploration of ligand pairs without serial computational bottlenecks.

Completion of all transitions and analyses required approximately 15 GB of storage per edge when using three replicas per environment.

### D. Length of the production

evaluate the influence of production length on the accuracy of the non-equilibrium transitions, we tested three simulation durations: 5, 20, and 50 ns per replica. At a temperature of 298 K, the analysis was performed for 100 evenly spaced transitions, each lasting 200 ps. Both protocols were applied to a representative subset of ligand pairs in the three environments (water, membrane, and MEP). The resulting free energy estimates showed no statistically significant differences between the three lengths, with no considerable difference between the mean value of one production and the other, except a 2.6 kcal·mol^−1^ difference in the 11a→11b edge between the production of 5 and 50 ns (Fig. S1, S2, S3). Consequently, all subsequent production simulations were performed using a 20 ns trajectory, which provided equivalent accuracy at substantially reduced computational cost while maintaining an acceptable error.

### E. Number de transitions

To assess the influence of the number of non-equilibrium transitions on the calculated free energies, we performed tests using 10, 20, 25, 33, 50, 80, and 100 evenly spaced transitions per edge of 200 ps, while maintaining a temperature of 298 K and a production run of 20 ns (the first transition starting from 5 ns). The results, summarized in Tab. S2, show that increasing the number of transitions beyond 80 had no significant impact on the predicted relative free energies (maximum deviations ≤0.05 kcal · mol^−1^ across all tested edges). However, increasing the number of transitions reduces the error. Based on this analysis, subsequent production simulations were performed using 50 uniformly spaced transitions, minimizing the number of transitions while maintaining a reasonable error.

### E. Length of the transitions

To evaluate the effect of transition length on the accuracy and convergence of the non-equilibrium work distributions, we tested 50 ps, 100 ps, and 200 ps transition durations for representative ligand pairs in all three environments (water, membrane, and MEP) and using three replicas. While maintaining a temperature of 298 K and a production run of 20 ns, as shown in Supplementary Fig. S1, S2, S3, transitions of 50 ps produced inconsistent RBFE estimates and showed insufficient overlap between forward and reverse work distributions, indicating inadequate sampling and poor convergence. In contrast, 100 ps and 200 ps transitions yielded highly consistent RBFEs, differing by less than 0.2 kcal·mol^−1^ and exhibiting reduced statistical uncertainty. Therefore, a transition length of 100 ps was chosen as the default in NEMAT, providing an optimal balance between accuracy and computational efficiency.

In conclusion, the benchmark was conducted using a production dynamic of 20 ns and 50 transitions evenly spaced, lasting 100 ps. These values are the defaults when using NEMAT, but can be easily modified. However, it is reasonable to start testing your system from these values.

**Table 1.**
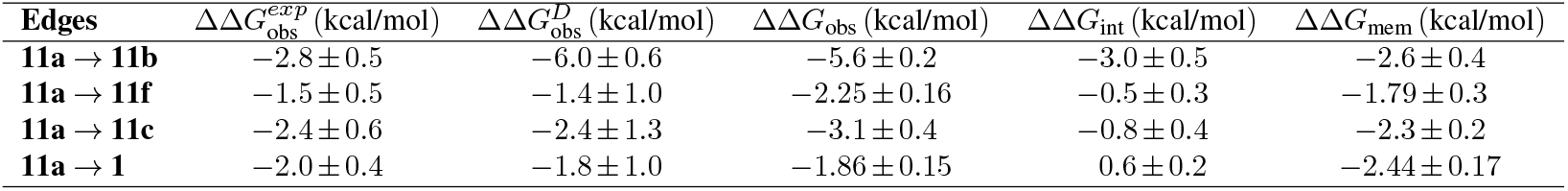
Comparison of Results. Table presenting the results obtained experimentally by Chao et al. (48) 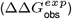, the results obtained by Dickson et al. without using Arsenic (the ones that come from adding ΔΔ*G*_int_ and 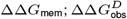) (26), and by NEMAT for the 11a group.

**Table 2.** Comparison of Results. Table presenting the results obtained experimentally by Chao et al. (48) 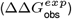, the results obtained by Dickson et al. without using Arsenic (the ones that come from adding ΔΔ*G*_int_ and 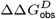) (26), and by NEMAT for the 6a group.

| Edges | $\Delta\Delta G_{obs}^{exp}$ (kcal/mol) | $\Delta\Delta G_{obs}^D$ (kcal/mol) | $\Delta\Delta G_{obs}$ (kcal/mol) | $\Delta\Delta G_{int}$ (kcal/mol) | $\Delta\Delta G_{mem}$ (kcal/mol) |
| --- | --- | --- | --- | --- | --- |
| 6a → 6i | $-1.0 \pm 0.4$ | $-2.7 \pm 0.3$ | $-2.7 \pm 0.2$ | $0.8 \pm 0.3$ | $-3.5 \pm 0.3$ |
| 6a → 6f | $-0.1 \pm 0.5$ | $-1.5 \pm 0.1$ | $0.72 \pm 0.08$ | $-0.5 \pm 0.2$ | $1.2 \pm 0.3$ |
| 6a → 6h | $-0.8 \pm 0.7$ | $-0.8 \pm 0.4$ | $-0.69 \pm 0.12$ | $1.03 \pm 0.18$ | $-1.72 \pm 0.15$ |
| 6a → 6m | $0.6 \pm 0.5$ | $-0.7 \pm 1.2$ | $2.8 \pm 0.3$ | $2.8 \pm 0.3$ | $0.0 \pm 0.3$ |
| 6a → 6g | $-0.5 \pm 0.6$ | $-1.6 \pm 0.4$ | $-3.3 \pm 0.3$ | $-2.9 \pm 0.5$ | $-0.4 \pm 0.5$ |
| 6a → 6l | $-1.2 \pm 0.7$ | $-0.0 \pm 0.5$ | $1.24 \pm 0.16$ | $-0.9 \pm 0.2$ | $0.3 \pm 0.2$ |
| 6a → 6j | $0.4 \pm 0.7$ | $-3.1 \pm 0.4$ | $-0.1 \pm 0.3$ | $1.0 \pm 0.3$ | $-1.1 \pm 0.2$ |
| 6a → 6n | $0.7 \pm 0.5$ | $-0.3 \pm 0.7$ | $2.48 \pm 0.13$ | $0.38 \pm 0.17$ | $2.11 \pm 0.14$ |

## Conclusions

In this work, we introduced NEMAT, the first automated pipeline for performing non-equilibrium alchemical free-energy calculations in explicit water, membrane, and membrane-embedded protein environments. By combining pmx-based hybrid topology generation with GPU-accelerated GROMACS simulations, NEMAT provides a fully reproducible workflow that decomposes observed affinities (ΔΔ*G*_obs_) into membrane partitioning (ΔΔ*G*_mem_) and receptor-interaction (ΔΔ*G*_int_) components. Applied to the P2Y_1_ receptor and a series of BPTU analogues, NEMAT reproduced experimental trends with accuracy comparable to established equilibrium approaches while maintaining excellent computational efficiency and parallel scalability.

Although NEMAT offers a practical and broadly applicable framework, several challenges remain. The optimal NEQ-FEP settings can vary by system, and parameterization with current force fields may limit performance for flexible or chemically complex ligands. In addition, heterogeneous membrane compositions and slow lipid or protein motions can introduce sampling difficulties that the present implementation does not fully address.

These limitations highlight opportunities for future development, including broader force-field support, enhanced sampling strategies, and improved handling of complex or asymmetric membranes. Such extensions would increase robustness and expand the range of systems that can be modeled reliably.

In summary, NEMAT fills a critical gap in the free-energy landscape by enabling fast, reproducible, and membrane-aware non-equilibrium calculations for ligand binding. Its ability to separate membrane and protein contributions provides new mechanistic insight into ligand selectivity at lipid-facing pockets, and continued development will help establish NEQ-FEP as a routine tool in membrane-protein drug discovery.

## Supporting information

Supplementary Information including methods details and supplementary figures

## ACKNOWLEDGEMENTS

The authors would like to acknowledge fruitful discussions with Berta Bori-Bru and Raul Santiago. This work was supported by the Ministerio de Ciencia, Innovación y Universidades under project PID2022-138040OB-I00. R.C. belongs to a consolidated research group (Grup de Recerca) of the Generalitat de Catalunya and has support from the Departament d’Universitats, Recerca i Societat de la Informació de la Generalitat de Catalunya (Reference: 2021 SGR 00476). The calculations described in this work were carried out at the Centre de Supercomputació de Catalunya (CSUC).

