## Supplementary Information including methods details and supplementary figures for "NEMAT: An Automated Non-Equilibrium Free-Energy Framework for Predicting Ligand Affinity in Membrane Proteins"

In membrane-embedded protein systems, ligand binding is further complicated by the physicochemical role of the lipid bilayer, which introduces additional kinetic and thermodynamic factors. For example, before a ligand can bind to a membrane protein's site, especially an extra-helical or lipid-exposed site, it must first partition into the membrane (26). Furthermore, membrane partitioning increases the local concentration of the ligand near the receptor, enhancing the observed binding affinity, even if the ligand's true interaction with the binding site is weak (31).

Another factor that complicates the affinity of a membrane-embedded receptor is that the ligand's affinity to the membrane may be similar to its affinity to the receptor, leading to a misperception of the true affinities with the protein. As stated in (28–30), if there is binding when a membrane is involved, there is a clear correlation between drug concentration and the observed binding affinity. This means that the aggregated association rate constant of the ligand ($K_{aa}$) is concentration dependent:

$$K_{aa} = [L] \cdot K_a + K_d \qquad \textbf{(4)}$$

Where $K_a$ is the association rate constant and $K_d$ is the dissociation rate constant, which reflects the true stability of the binding. Then, knowing that association is proportional to the concentration of the ligand, but $k_d$ is not, Sykes *et al.* (29) observed a direct correlation between $k_a$ and a new constant which reflects the membrane binding $K_{\text{mem}}$. Then we can obtain an estimation of the "real" binding of the ligand ($K_{\text{int}}$) with:

$$pK_{\text{int}} = pK_{\text{obs}} - pK_{\text{mem}} \qquad \textbf{(5)}$$

If alchemical transformations are performed in water, in the membrane, and the protein + membrane system, a thermodynamic cycle can be created such that the relations among relative free energies, based on EQ. 5, become:

$$\Delta G_{\text{obs}} = \Delta G_{\text{mem}} + \Delta G_{\text{int}} \qquad \textbf{(6)}$$

Such that $\Delta G_{\text{obs}}$, $\Delta G_{\text{mem}}$ and $\Delta G_{\text{int}}$ are the relative free energy between the protein + membrane system and water, between the membrane and the protein + membrane system and between the membrane and water system respectively as can be observed in Fig. 2.

The GROMACS parameters specific for FEP transitions of 100 ps are presented in TAB. S1.

| Prameter | Value |
|---:|---|
| `free-energy` | `yes` |
| `init-lambda` | `0` |
| `delta-lambda` | $2{\cdot}10^{-5}$ |
| `sc-alpha` | `0.3` |
| `sc-sigma` | `0.25` |
| `sc-power` | `1` |
| `sc-coul` | `yes` |

**Table S1. Parameters for FEP used for transitions.** The parameter *init-lambda* (0 or 1) indicates which ligand the transformation starts from. *delta-lambda* must be $1/nsteps$ and negative if *init-lambda* is 1.

### A. Testing numbers. About the length of the production and the transitions.

About the number of transitions

| Edge | 100 ti | 80 ti | 50 ti | 33 ti | 25 ti | 20 ti | 10 ti |
|---|---|---|---|---|---|---|---|
| 11a → 11b | $-5.98 \pm 0.13$ | $-6.0 \pm 0.2$ | $-5.9 \pm 0.3$ | $-5.9 \pm 0.3$ | $-5.9 \pm 0.4$ | $-5.4 \pm 0.4$ | $-5.3 \pm 0.5$ |
| 11a → 11f | $-2.00 \pm 0.11$ | $-2.1 \pm 0.3$ | $-1.9 \pm 0.4$ | $-2.0 \pm 0.4$ | $-2.0 \pm 0.4$ | $-2.1 \pm 0.6$ | $-1.6 \pm 0.7$ |
| 11a → 11c | $-3.42 \pm 0.14$ | $-3.4 \pm 0.2$ | $-3.6 \pm 0.3$ | $-3.5 \pm 0.4$ | $-3.5 \pm 0.4$ | $-4.1 \pm 0.4$ | $-4.3 \pm 0.4$ |
| 11a → 1 | $-1.39 \pm 0.11$ | $-1.34 \pm 0.10$ | $-1.31 \pm 0.13$ | $-1.35 \pm 0.15$ | $-1.23 \pm 0.13$ | $-1.4 \pm 0.2$ | $-1.2 \pm 0.3$ |

**Table S2. Results obtained using different numbers of frames.** Values obtained from a single replica for the $\Delta\Delta G_{\text{obs}}$ of the 11a group. The values correspond to a production of 20 ns (starting transitions from 5 ns) and transitions lasting 200 ps. The transitions are evenly spaced.

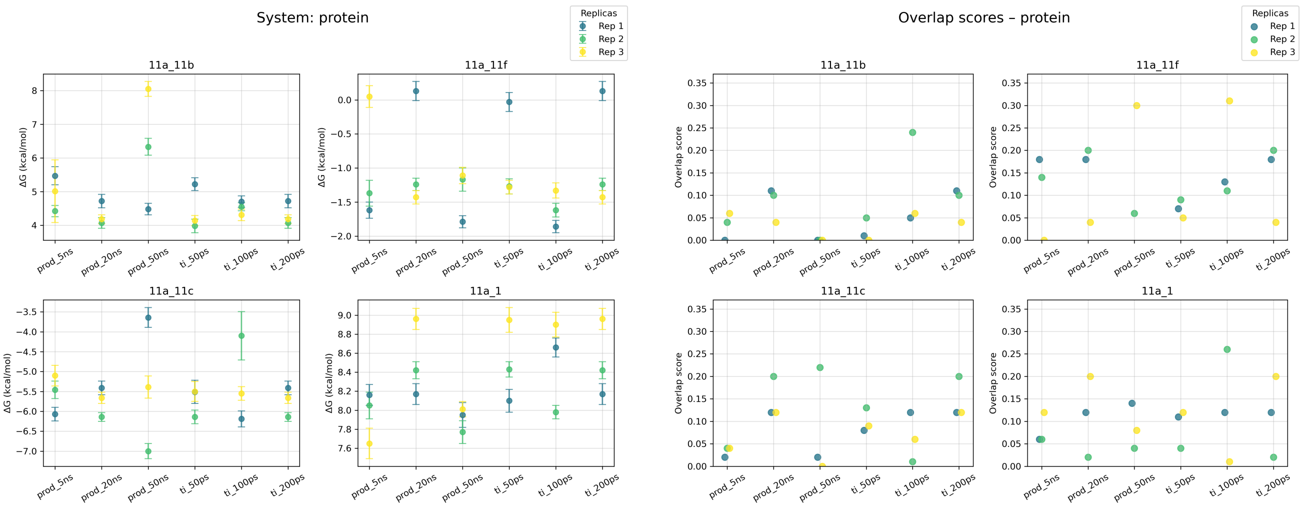


**(a)** Computed $\Delta G_p$ (Fig. 1) for the 11a ligand subsystem. **(b)** Overlap between forward and backwards trajectories of the 11a ligand subsystem.

**Fig. S1. Tests for the MEP system of overlap and results.** The results of 3 replicas are plotted.

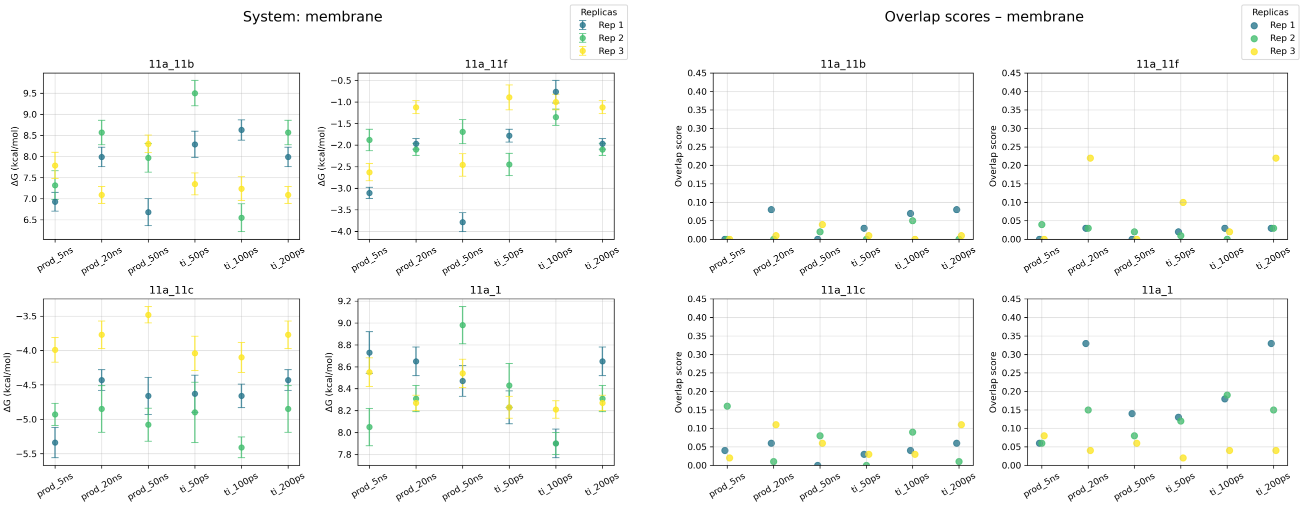


**(a)** Computed $\Delta G_m$ (Fig. 1) for the 11a ligand subsystem. **(b)** Overlap between forward and backwards trajectories of the 11a ligand subsystem.

**Fig. S2. Tests for the membrane system of overlap and results.** The results of 3 replicas are plotted.

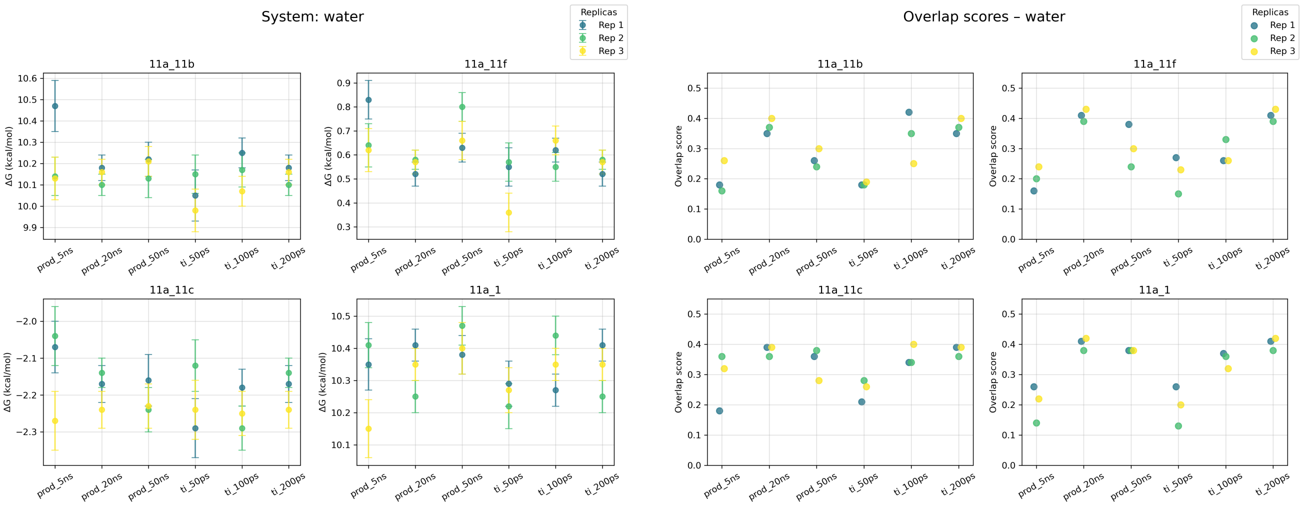


**(a)** Computed $\Delta G_w$ (Fig. 1) for the 11a ligand subsystem. **(b)** Overlap between forward and backwards trajectories of the 11a ligand subsystem.

**Fig. S3. Tests for the water system of overlap and results.** The results of 3 replicas are plotted.

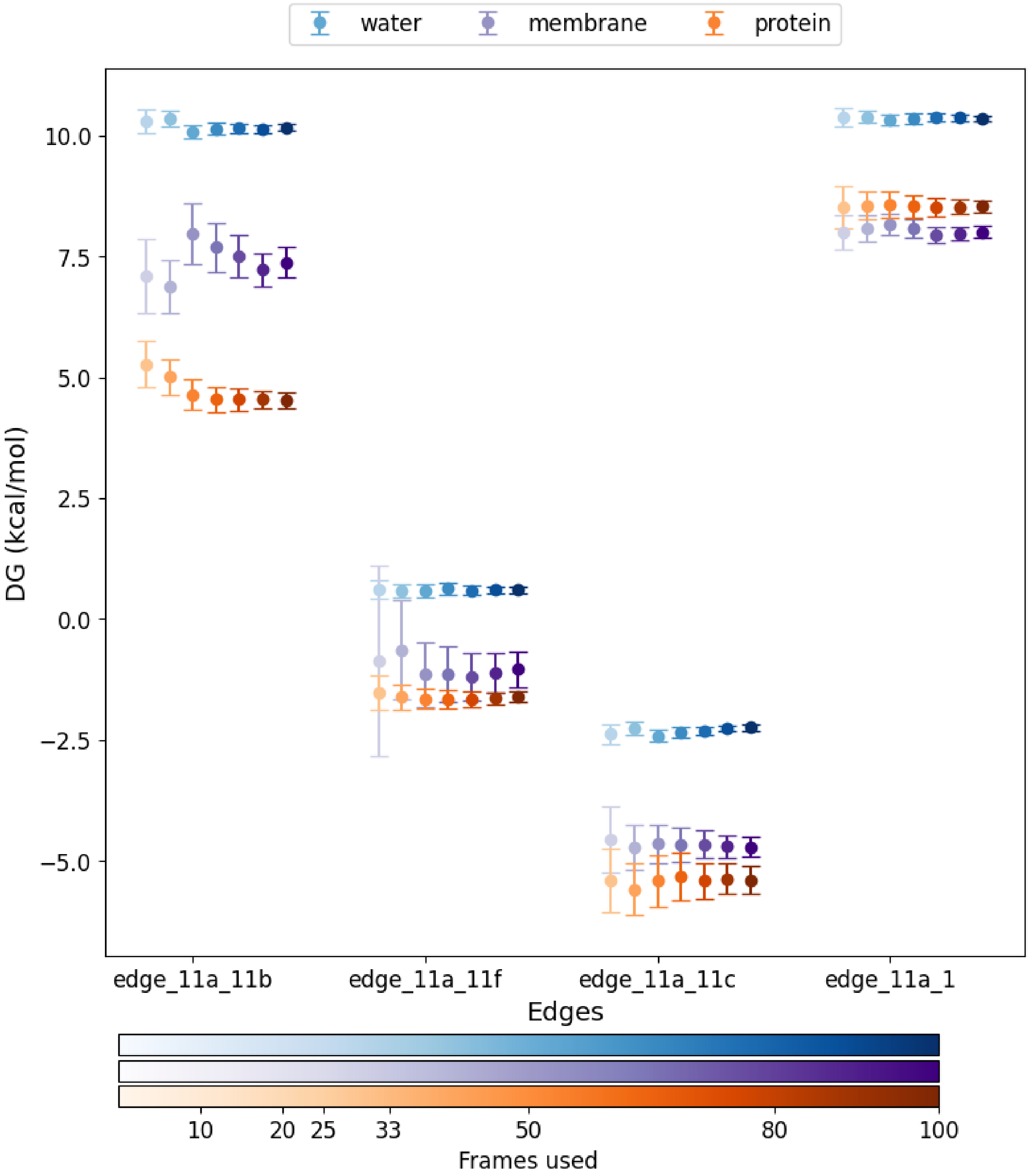


**Fig. S4. Example of the effect of using fewer transitions.** Reducing the number of transitions drastically reduces the sampling of uneven behavior and increases its associated error, potentially leading to different results.

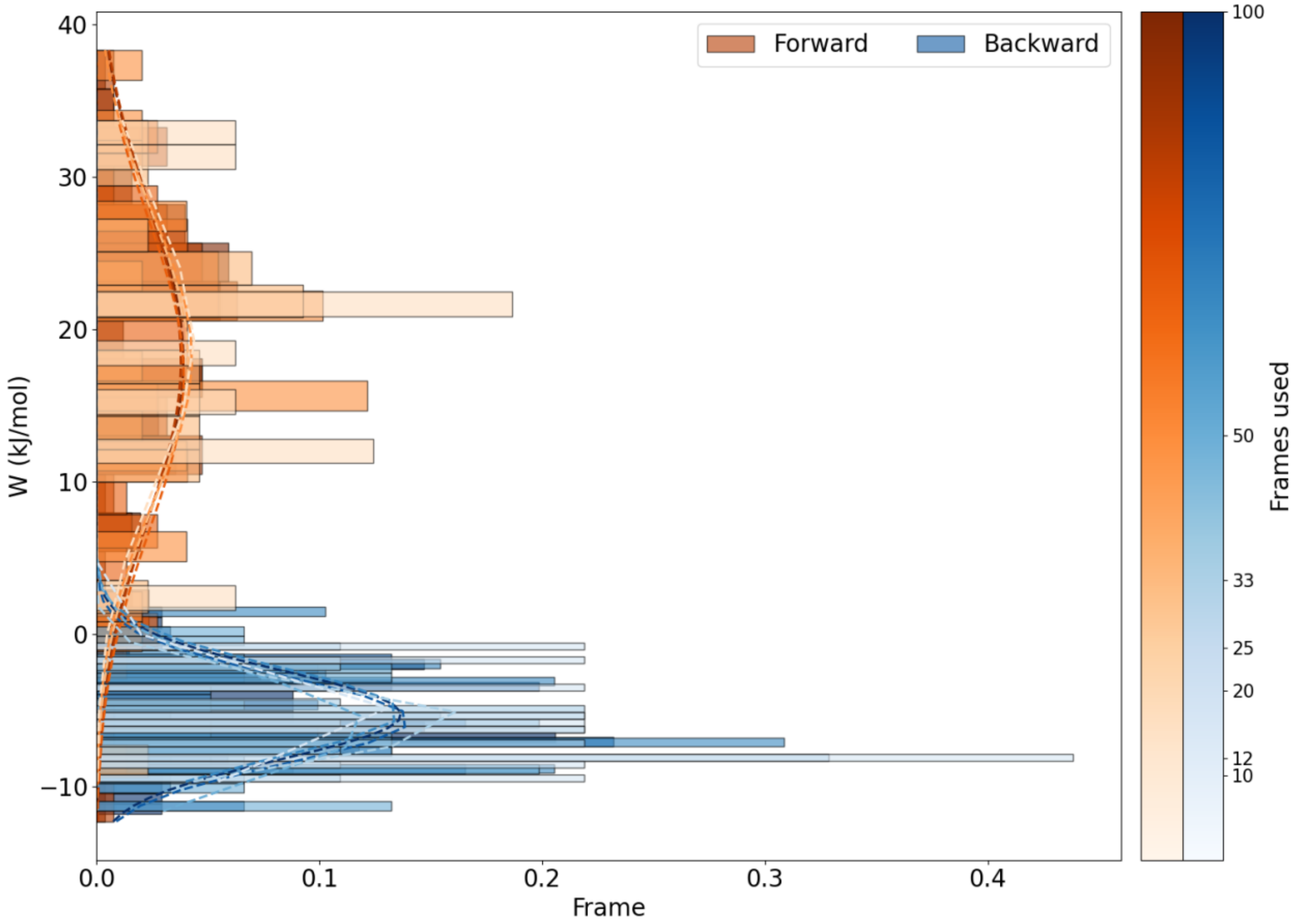


**Fig. S5. Overlap between forward and backwards trajectories of the 11a ligand with the membrane subsystem.** In the example, we can observe the distribution change when reducing the number of transitions. There is a small shift in the intersection point of the distributions.

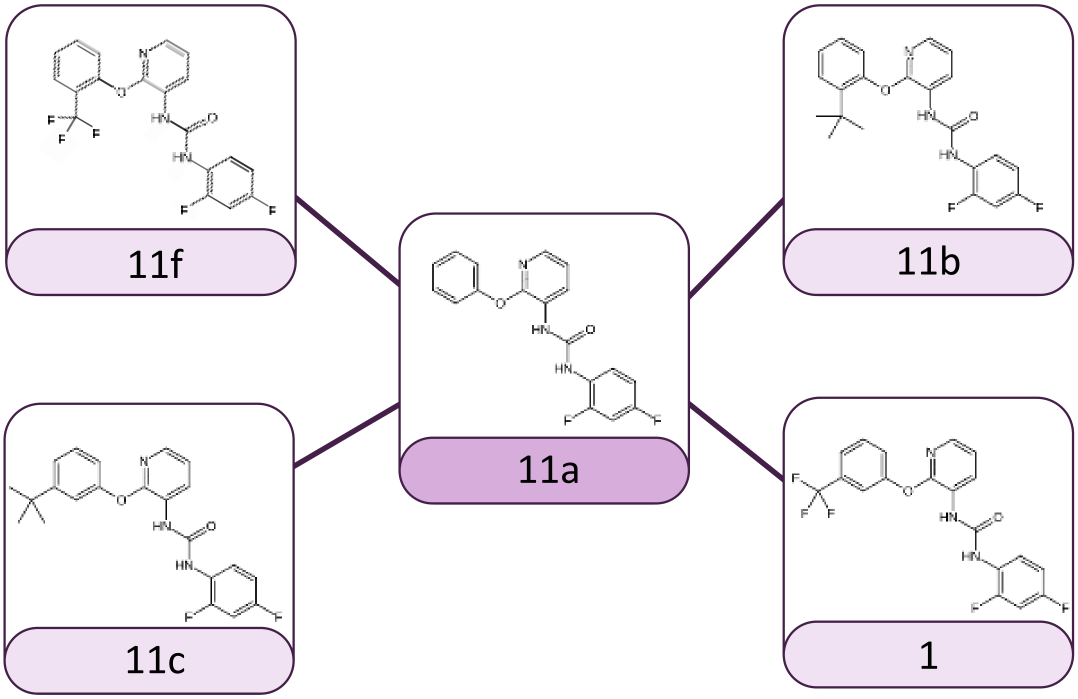


**Fig. S6. Start map centered on 11a for the BPTU analogues.**

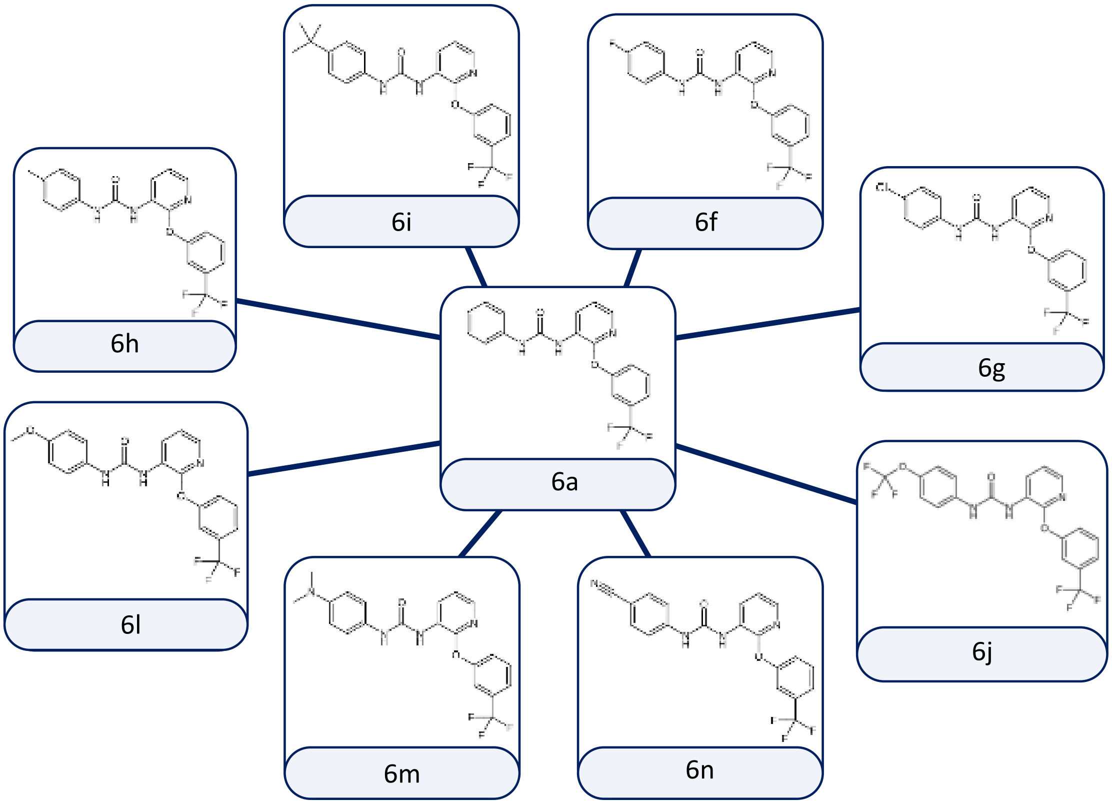


**Fig. S7. Start map centered on 6a for the BPTU analogues.**
